# Characterization of the Cellular Immune Response to Group B Streptococcal Vaginal Colonization

**DOI:** 10.1101/2025.01.29.635275

**Authors:** Brady L. Spencer, Dustin T. Nguyen, Stephanie M. Marroquin, Laurent Gapin, Rebecca L. O’Brien, Kelly S. Doran

## Abstract

**Introduction:** Group B *Streptococcus* (GBS) asymptomatic colonizes the female genital tract (FGT) but can contribute to adverse pregnancy outcomes including pre-term birth, chorioamnionitis, and neonatal infection. We previously observed that GBS elicits FGT cytokine responses, including IL-17, during murine vaginal colonization; yet the anti-GBS cellular immune response during colonization remained unknown. We hypothesized that GBS may induce cellular immunity, resulting in FGT clearance.

**Methods:** Herein, we utilize depleting antibodies and knockout mice and performed flow cytometry to investigate cellular immunes responses during GBS colonization.

**Results:** We found that neutrophils (effectors of the IL-17 response) are important for GBS mucosal control as neutrophil depletion promoted increased GBS burdens in FGT tissues. Flow cytometric analysis of immune populations in the vagina, cervix, and uterus revealed, however, that GBS colonization did not induce a marked increase in FGT CD45+ immune cells. We also found that that Vγ6+ γδ T cells comprise a primary source of FGT IL-17. Finally, using knockout mice, we observed that IL-17-producing γδ T cells are important for the control of GBS in the FGT during murine colonization.

**Conclusions:** Taken together, this work characterizes FGT cellular immunity and suggests that GBS colonization does not elicit a significant immune response, which may be a bacterial directed adaptive outcome. However, certain FGT immune cells, such as neutrophils and ɣδ T cells, contribute to host defense and control of GBS colonization.

## Introduction

Group B *Streptococcus* (GBS), or *Streptococcus agalactiae,* is a Gram-positive, β-hemolytic bacterium that asymptomatically colonizes the gastrointestinal tract and the female genital tract (FGT) of 10-30% of healthy adults in the United States [1]. GBS can cause infection in immunocompromised adults [2], and it presents problems in pregnancy as ascension of vaginally colonizing GBS to the uterus can cause complications including chorioamnionitis and preterm births [3, 4]. Further, GBS neonatal early-onset disease (EOD) results from maternal transmission to the neonate *in utero* or during birth, presenting within 7 days after birth as pneumonia, bacteremia, and in severe cases, meningitis [5]. GBS is a leading cause of neonatal infection and causes 2-3% mortality of term newborns and up to 30% of preterm newborns who develop GBS disease [6–8]. Despite this, no GBS vaccine is currently available, and antibiotic prophylaxis is the primary treatment for pregnant women testing positive for GBS recto-vaginal swab at 37 weeks [4]. Even with this intervention, GBS remains a global health issue as the primary risk factor for neonatal EOD is maternal vaginal colonization with GBS [4]. Thus, understanding GBS-host interactions in the mucosal environment of the FGT is needed to develop new therapies.

Similar to other mucosal barriers, the FGT contains numerous defenses to prevent colonization and invasion of pathogens, such as physical epithelial and mucus barriers, chemical defenses (antimicrobial peptides, low pH), as well as the vaginal microbiota [9, 10]. Soluble and cellular immune responses within the FGT can also be initiated to control infection [11, 12]. While immune responses to viral and fungal pathogens have been characterized in the FGT, less work has been done to characterize cellular immune responses to vaginal pathobionts and opportunistic pathogens in this niche.

As an opportunistic pathogen, GBS encodes numerous virulence factors that allow it to persist in the mucosa and ascend to the uterus despite these host defenses, including adhesins and invasins (BspC, Srr proteins, pili) [13–15], a β-hemolysin/cytolysin [16–18], a hyaluronidase [19], capsular polysaccharide [20], and a type VII secretion system [21, 22]. While these factors have individually been shown to promote GBS colonization and/or ascending infection, previous work from our laboratory found that GBS clinical isolates exhibit a range of murine vaginal persistence over time [23] and a single GBS determinant responsible for this differential clearance has yet to be identified. Previous work from our laboratory and others have also profiled cytokine responses in response to various GBS isolates, assessing both *in vitro* cytokine analysis from GBS infection of vaginal epithelium as well as cytokine induction within GBS-colonized FGT tissue homogenates, with some isolates eliciting more cytokine induction than others [23, 24]. Based on this, we hypothesized that GBS strains may elicit cellular immune responses during colonization and ascension to the uterus, which result in their clearance from the FGT. In this study, we profiled the cellular immune responses to persistent-colonizing serotype V GBS isolate CJB111, and to serotype III GBS isolate COH1 in the vagina, cervix, and uterus, and show the importance of specific immune populations in controlling GBS during FGT colonization.

## Methods

### Mice

CD-1 (outbred Swiss) mice were obtained from Charles River Laboratories. C57BL/6J mice were originally obtained from Jackson Laboratory and bred at National Jewish Health (Denver, CO) and then University of Colorado-Anschutz (Aurora, CO). TCR-Vγ4^-/-^/Vγ6^-/-^ mice in the C57BL/6J genetic background were originally derived from TCR-Vγ4^-/-^/Vγ6^-/-^ mouse line generated by Dr. Koichi Ikuta (Kyoto University, Kyoto, Japan).

### Bacterial strains and growth

GBS isolates COH1 [25] and CJB111 [26] were grown statically in Todd-Hewitt Broth (THB; Research Products International, RPI Catalog# T47500) at 37°C. For vaginal colonization experiments, bacteria were grown to mid-logarithmic phase (OD_600_ of approximately 0.4) and normalized in PBS to 1x 10^9^ CFU/mL.

### Neutrophil depletion in mice

Murine neutrophil depletion protocols were adapted from [27]. Depleting α-Ly6G (BioXCell, clone 18A) and IgG2a isotype control (BioXCell, clone 2A3) antibodies were diluted in buffered reagent diluent to a concentration of 2 mg/mL and animals were intraperitoneally injected with 100 μL (dose of 200 μg) the day prior to vaginal bacterial inoculation. A second dose was administered intraperitoneally three days later at day 2 post-GBS inoculation.

### Murine model of vaginal colonization

GBS vaginal colonization and tissue invasion were assessed using a previously described murine model of vaginal persistence and ascending infection [28]. 8-week-old female CD-1 outbred mice or 8-16-week-old C57BL/6J and TCR-Vγ4^-/-^/Vγ6^-/-^ mice in the C57BL/6J genetic background were synced with β-estradiol at day −1 and inoculated with 1 × 10^7^ GBS in 10 µL of phosphate buffered saline (PBS) on day 0. Post-inoculation, mice were vaginally lavaged with PBS, and the samples were serially diluted and plated for CFU quantification to determine bacterial persistence on differential and selective GBS CHROMagar (catalog# SB282(B)). At experimental time points, mice were euthanized, and FGT tissues were harvested. In experiments using knockout mice or neutrophil depletion, tissues were homogenized and then serially diluted and plated on CHROMagar for CFU enumeration. In experiments assessing immune responses, tissues were processed as described below, serially diluted, and plated on CHROMagar for CFU enumeration. Bacterial counts were normalized to tissue weight.

### Dissociation of FGT tissues for flow cytometry

To evaluate cellular immune responses in the FGT, following euthanasia, mice were perfused with PBS and heparin through left ventricle cardiac puncture. Vaginal, cervical, and uterine tissue were obtained from terminal harvest, minced with a razor blade, and suspended in 2.4 mL RPMI 1640 medium (RPMI; Corning). Digestion enzymes were obtained from Miltenyi Biotec Multi Tissue Dissociation Kit 1 and resuspended according to manufacturer instructions. 100 μL Enzyme D, 50 μL Enzyme R, 12.5 μL Enzyme A were added to each tissue sample in RPMI. Tissues were mechanically dissociated using gentleMACS™ C Tubes (Miltenyi Biotec; protocol Multi H). Tissues suspensions were placed in a rocking shaker at 37ᵒC for 1 hour, with an additional 30 second mechanical dissociation half-way through the incubation. The digestion mixture was strained through a 70 μm filter, and the single cells were spun down at 300 x g for 5 minutes and washed with RPMI. The cell pellet was resuspended in Ammonium-Chloride-Potassium red blood cell lysis buffer (150 mM NH_4_Cl, 10 mM KHCO_3_, 0.1 mM Na_2_EDTA) for 1 minute at room temperature and diluted in RPMI. Following centrifugation, the cell pellet was resuspended in MACS staining buffer (1X PBS, 2 mM EDTA, 0.5% BSA) to generate single cell suspensions.

### Antibody staining of single cell suspension for flow cytometry analysis

Fixable Viability Dye eFluor™ 506 (ThermoFisher; 1:800 dilution) in PBS was added to single cell suspensions and allowed to incubate for 15 minutes at room temperature. The innate flow cytometric panel was adapted from [29] and a T cell flow cytometric panel was developed in the present study. Conjugated antibodies listed in **Supplemental Table 1** were then added to single cell suspensions along with anti-mouse CD16/32 (ThermoFisher, clone 2.4G2) to block mouse Fc receptors for 30 minutes at room temperature. Stained cells were then washed and prepared for fixation and permeabilization using Foxp3/Transcription Factor Staining Buffer Set (eBioscience) according to the manufacturer’s protocol. Prior to running samples on the flow cytometer, conjugated antibodies for intracellular targets were added to cells in permeabilization buffer for 30 minutes at room temperature. Cells were washed and resuspended and analyzed on the CytoFlex (Beckman Coulter) and LSRFortessa (BD Biosciences) flow cytometers. Data were analyzed with BD FlowJo software version 10.10.0. Single-stained beads (Versacomp Antibody Capture Bead Kit; Beckman Coulter) were used as compensation controls.

A detailed gating strategy was established based on single stain and fluorescence minus one controls (**SFig 2a, b**). In this study, we found that cells from a subset of mice were not recognized by our anti-MHCII antibody (clone M5/114 15.2), which reacts with a polymorphic determinant shared by the I-A^b^, I-A^d^, I-A^q^, I-E^d^, and I-E^k^ MHC class II alloantigens, but does not react with I-A^f^, I-A^k^, or I-A^s^ MHC class II alloantigens. We expect that these mice carry other MHC class II haplotypes, but this would need to be experimentally shown. Because MHCII staining impacts our innate gating strategy, mice exhibiting negative MHCII staining were excluded from our innate immune analysis. We similarly observed that cells from some CD-1 outbred mice did not react with our anti-Ly6C antibody, and the reason for this remains unclear. Because of this we did not use Ly6C expression to define populations within our gating strategy.

### Stimulation assays

Single cell suspensions of FGT tissues were prepared for flow cytometry as previously stated. The final pellet was resuspended in 2 mL of cell stimulation media: RPMI with L-glutamine, supplemented with 10% FBS, 1X non-essential amino acids, 1mM sodium pyruvate, 10 mM HEPES (Gibco). Cells were stimulated with 1X eBioscience Cell Stimulation Cocktail (Thermo, 00-4970-93) containing PMA and ionomycin for one hour at 37C. GolgiStop™ Protein Transport Inhibitor (BD Biosciences, 554724) was added at a 6.67uL/mL media concentration at this time and cells were incubated for an additional 4 hours at 37ᵒC, for a total stimulation time of 5 hours. Cells were washed twice with MACS staining buffer and flow cytometric staining progress as described above with the addition of IL-17A-PE-Cy7 (Invitrogen, [eBio1787], 25-7177-80) antibody staining following fixation/permeabilization. In some assays, a PE-conjugated antibody to the Vγ6 TCR was included (Clone 1C10-1F7, BD Pharmingen, Catalog # 570217) to specifically identify Vγ6+ ɣδ T cells [30].

### Statistical Analysis

Data were analyzed using PRISM9 (GraphPad Software). Data are depicted as mean ± SEM or median where indicated. P-values are denoted by *(P<0.05), **(P<0.01), ***(P<0.001), and ****(P<0.0001).

## Results

### Neutrophils control GBS burdens during FGT colonization

Previous work from our laboratory identified elevated IL-17 levels during CJB111 FGT colonization and found that proportions of IL-17^+^ immune populations were higher in mice that had cleared GBS [23]. However, the downstream implications of IL-17 signaling were not investigated. IL-17 family cytokines are known to stimulate production of chemokines such as G-CSF and IL-8, resulting in the recruitment of neutrophils [31]. In this study, to determine if neutrophils contribute to the control of GBS FGT colonization and uterine ascension, we performed systemic neutrophil depletion using an anti-Ly6G antibody. CD-1 outbred mice were administered IP injections of anti-Ly6G or an isotype control IgG2a the day prior to GBS inoculation and two days post-GBS inoculation as shown in **Figure 1a** and neutrophil depletion was confirmed in FGT tissues at day 4 by flow cytometry (**SFig. 1**). To determine the functional impact of neutrophils on bacterial burden during colonization, we assessed GBS CFU in FGT tissues of mice in the presence or absence of neutrophil depletion at 4 days post-inoculation. We found that neutrophil-depleted mice exhibited higher burdens of GBS in the vagina, cervix, and uterus compared to non-depleted isotype-injected controls (**Fig. 1b-d**). This was also observed in the vagina and uterus using another GBS strain, COH1 (**Fig. 1e-g**). These data suggest that neutrophils may help to control GBS burdens during colonization.

**Figure 1.**
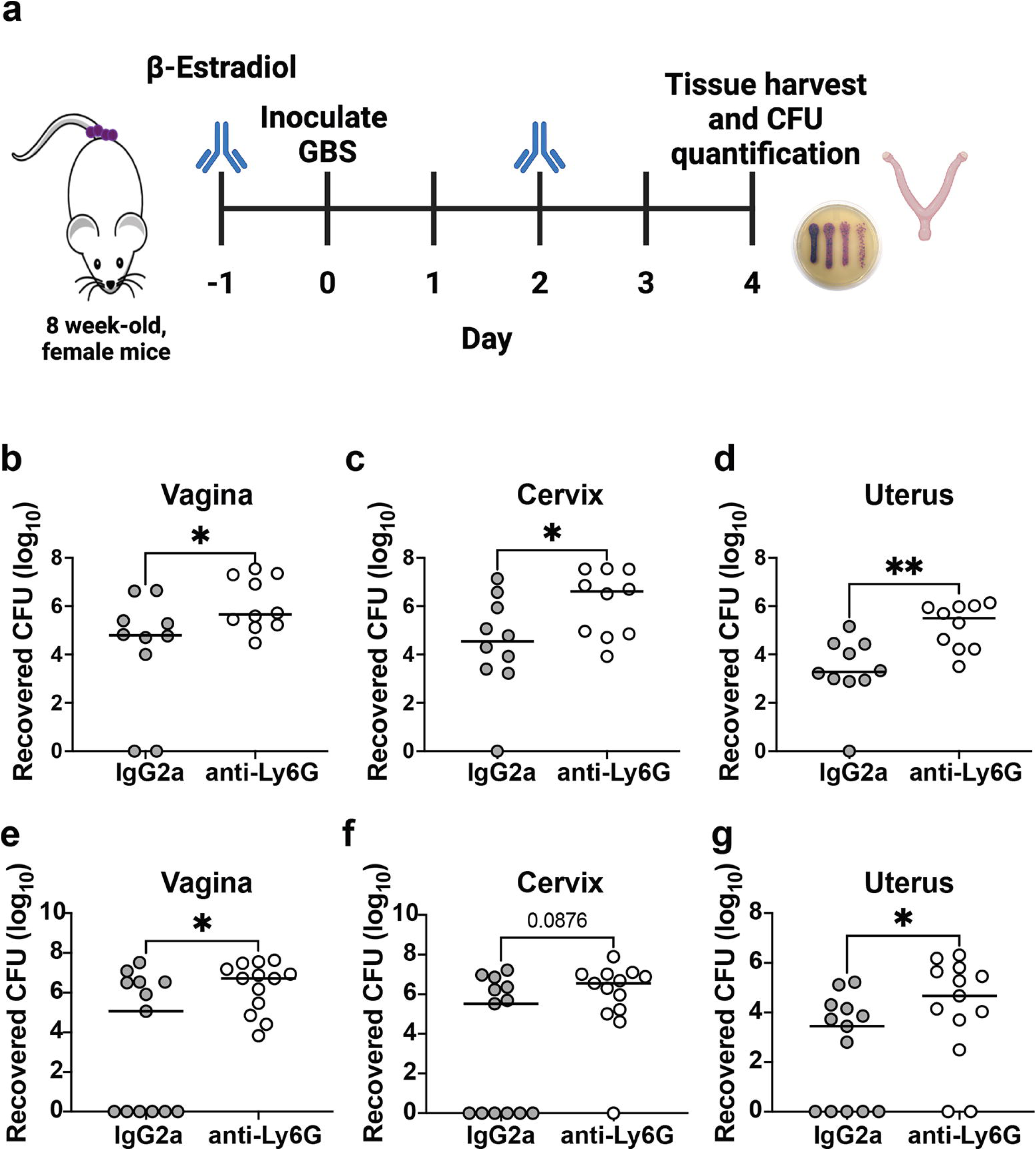
Systemic depletion of neutrophils results in increased GBS burdens in FGT tissues. **a**) Schematic of systemic neutrophil depletion in mice during vaginal colonization. Mice were treated twice with anti-Ly6G or isotype control IgG2a (days −1 and 2) through intraperitoneal injection during vaginal colonization to systemically deplete neutrophils. Mice were colonized with GBS intravaginally on day 0 following β-estradiol injection at day −1 and were euthanized on day 4 for tissue harvest to enumerate CFU in FGT tissues. Recovered CFU counts from the **b, e**) vaginal **c, f**) cervical, and **d, g**) uterine tissue of CJB111 (**b-d**) or COH1-colonized mice (**e-g**). Each dot represents one mouse with n = 10 and 13/group in CJB111 and COH1 experiments, respectively. The bars in these plots show the median and statistics represent the Mann Whitney U test.

### Assessment of FGT immune populations during GBS colonization by flow cytometry

We next sought to evaluate changes in immune populations within FGT tissues during colonization using a previously published innate panel as well as a T cell panel we developed here to assess immune cells in FGT tissues (**SFig. 2a, b**). This innate panel allows for identification of various populations including neutrophils (CD11b^+^ Ly6G^+^), SSClo/MHCII^-^ (including NK cells and monocytes), dendritic cells (CD11c^+^ MHCII^+^), macrophages (CD11b^+^ F4/80^+^), as well as a CD24+ population (likely including eosinophils; CD11b^+^ CD24^+^ Ly6G^-^ F4/80^-^) [29]. Additionally, the T cell panel allows for identification of ɣδ T cells, and TCRβ^+^ T cells, including CD4^+^ and CD8^+^ T cells. To identify host cellular responses to GBS in the FGT, we vaginally inoculated CD-1 outbred mice with PBS or GBS isolate CJB111 and dissociated vaginal, cervical, and uterine tissues at 1, 4, 7, 14, and 28 days after GBS inoculation for flow cytometric analysis. We did not observe a significant difference in numbers of CD45^+^ immune cells (as a proportion of total single cells) in CJB111-colonized animals when compared to naïve animals over a time course of colonization (**Fig. 2a**). Similar results were observed with another GBS isolate, COH1 (**SFig. 3a**) [23].

**Figure 2.**
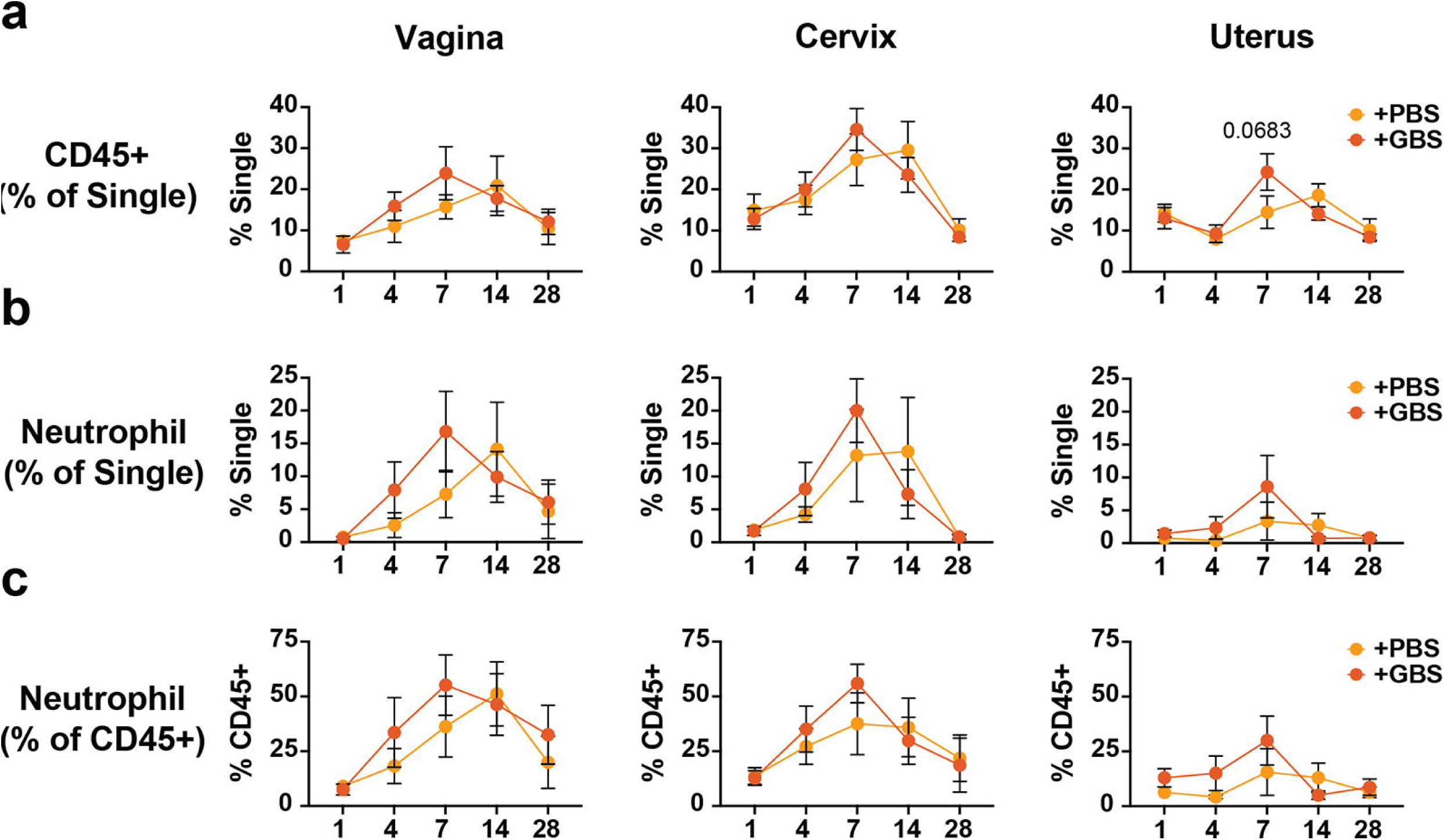
Total immune cells and neutrophils are not significantly altered over a time-course of GBS vaginal colonization. Line graphs depict **a**) CD45^+^ immune cells as a proportion of single cells and **b**) neutrophils (as a proportion of single cells) or **c**) as a proportion of CD45^+^ immune cells over a time-course of GBS colonization at days 1, 4, 7, 14, and 28. Yellow lines indicate PBS mock-colonized mice and orange lines indicate GBS-colonized mice. Each data point represents the mean and error bars indicate standard error of the mean. The data represents a total of n = 6-8 per timepoint and is combined across 2-3 independent experiments at each timepoint. Statistics represent two-way ANOVA with Šídák’s multiple comparisons test.

To evaluate the dynamics of the cellular immune response, we next examined specific cell innate and T cell populations over a time course of GBS colonization. We did not observe a significant increase in neutrophils in FGT tissues during GBS colonization compared to mock-colonized animals, although there was a slight but not significant trend for increased neutrophils in tissues of GBS-colonized mice at early timepoints (**Fig. 2b, c**). We similarly observed no significant differences in total number of neutrophils following colonization with another GBS isolate, COH1 (**SFig. 3b, c>**). Upon investigation of other innate and T cell populations, we consistently found no major changes in immune population proportions at the time points examined over the course of GBS colonization (**SFig. 4-5**). In fact, in some cases, such as for CD4^+^ and DN TCRβ+ T cell populations, GBS colonization appeared to slightly (but not significantly) lower immune cell percentages (**SFig. 5**). These data suggest that GBS may not dramatically alter the immune landscape of the FGT during cervicovaginal colonization of nonpregnant mice.

### Importance of IL-17+ immune populations to GBS clearance in the FGT

T cells are a major source of IL-17 [32, 33] and, in C57BL/6 mice it is known that γδ T cells are a primary source of IL-17 in the FGT [34]. To evaluate cellular sources of IL-17 in the FGT of CD-1 outbred mice, we harvested FGT tissues, generated single cell suspensions, and stimulated cells with PMA/ionomycin to elicit cytokine production (**Fig 3a**). Following stimulation, we found that γδ T cells also comprise the majority of FGT IL-17^+^ cells in CD-1 mice (**Fig 3b, c**). γδ T cells are among the first immune cells to take residence in the murine FGT during development (69). These FGT-resident γδ T cells express a Vγ6 T cell receptor (TCR) and are known to be IL-17 producing and to promote clearing of bacterial infections (70-72). To evaluate if FGT Vγ6+ γδ T cells are included in our observed IL-17 γδ T cell population, we repeated stimulation assays and concurrently stained for Vγ6. Indeed, we found that the majority of IL-17-producing γδ T cells in the FGT express the Vγ6 TCR (**Fig 3d**). Based on these results, we next hypothesized that resident Vγ6^+^ IL-17 producing ɣδ T cells may be important for controlling GBS burdens during vaginal colonization and ascension to the uterus. To investigate this, we utilized a previously characterized mouse line that lacks IL-17-producing Vγ4 and Vγ6 γδ T cells [35, 36]. Upon colonization of WT C57BL/6J or Vγ4^-/-^6^-/-^ mice with GBS, we found that GBS persisted longer in the vaginal lumen (**Fig 4a**), and within vaginal, cervical, and uterine tissue in mice lacking Vγ4^+^ or Vγ6^+^ γδ T cells (**Fig 4b-d**). These data indicate that IL-17-producing γδ T cells may be important for control of GBS colonization and subsequent infection.

**Figure 3.**
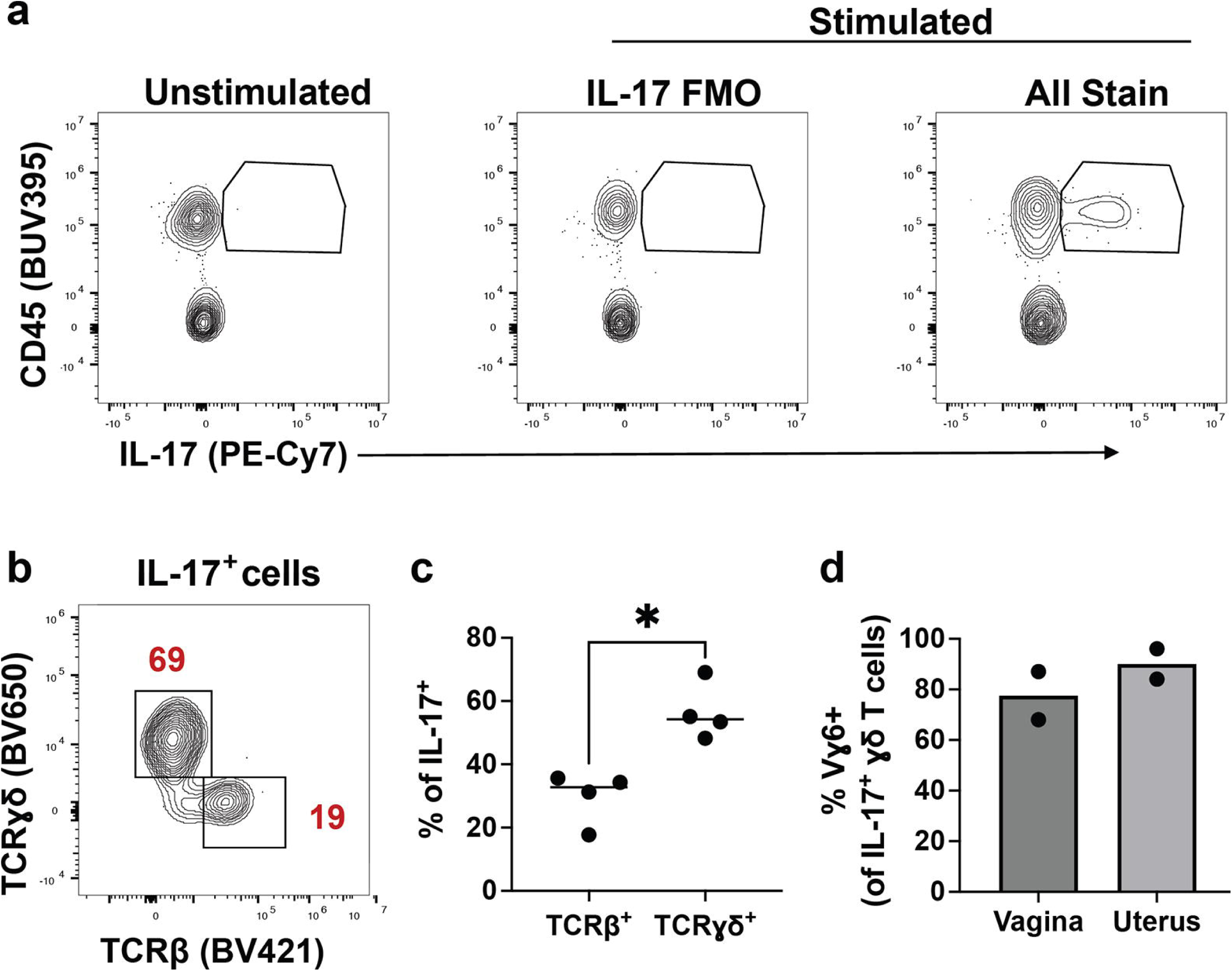
FGTɣδ T cells are a primary source of IL-17 production. **a**) Intracellular IL-17 production measured following five-hour FGT cell stimulation with PMA/ionomycin. IL-17^+^ gate drawn based on fluorescence minus one control. **b)** Representative contour plot of IL-17^+^ cells from stimulated pooled FGT single cell suspensions indicate that this population is comprised primarily by ɣδ T cells. Percentages are quantified in **c),** in which each dot represents one mouse (n = 4, across three independent experiments). Bars in these plots show the median, and statistics represent the Mann Whitney U test. **d)** Percentages of Vɣ6^+^ cells of IL-17^+^ ɣδ T cells indicate that this ɣδ T cell subset is the primary IL-17 producer in the FGT. Each dot represents one mouse (n = 2) and bars indicate the median.

**Figure 4.**
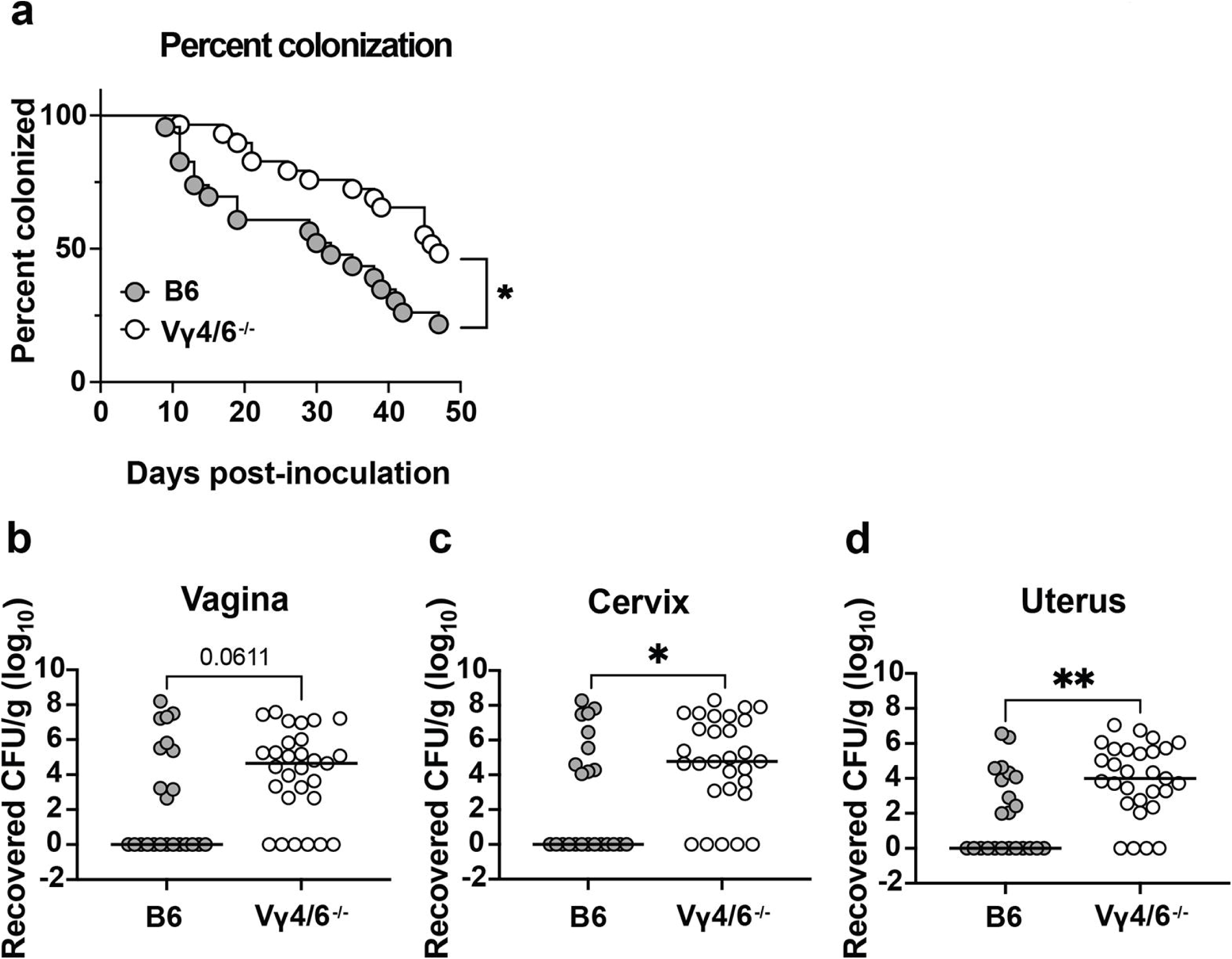
Loss of Vɣ4/6^+^ ɣδ T cells results in increased GBS burdens in female genital tract tissues. **a)** Percent colonization curves of WT C57BL/6J or Vɣ4/6^-/-^ female mice vaginally inoculated with CJB111 GBS. Statistics reflect the Log rank (Mantel-Cox) test. Recovered CFU counts from the **b**) vaginal **c**) cervical, and **d**) uterine tissue of colonized mice euthanized at day 47 post-inoculation. Each dot represents one mouse, bars in these plots show the median, and statistics represent the Mann Whitney U test. All graphs represent the combination of two independent experiments (n = 23 C57BL/6J, n = 29 Vɣ4/6^-/-^ total mice).

## Discussion

In this work, we have identified neutrophils as contributors to control of GBS during FGT colonization of non-pregnant CD-1 outbred mice. We further profiled the immune landscape of the FGT during GBS colonization and found that immune cell populations, including neutrophils, largely do not change over a time-course of colonization using two unique GBS isolates, with slight but not significantly increased trends observed in neutrophil populations at some early timepoints. We hypothesize that this may reflect an evolutionary adaptation by GBS to promote colonization in the FGT and to evade host clearance. IL-17 has been associated with GBS clearance previously and, in this study, we identified γδ T cells as major producers of IL-17 in the FGT of CD-1 outbred mice. We further show evidence for a contribution of IL-17-producing γδ T cells in controlling GBS during vaginal colonization and ascension to the uterus. These data highlight the commensal status of GBS in the FGT but also indicate that bacterial control by certain immune populations is possible and may be needed to keep colonization asymptomatic and in check.

Our flow cytometric panels enabled detection of a variety of innate populations, including neutrophils, SSClo/MHCII^-^ cells (including monocytes and NK cells), dendritic cells, and macrophages. We further assessed T cell populations, including ɣδ T cells, and TCRβ^+^ T cells (CD4^+^, CD8^+^, and double-negative (DN)). However, due to limitations in numbers of fluorophores in our conventional flow cytometric panels, we were not able to evaluate all immune populations. For example, while markers for mast cells were not included in our panel, mast cells have been shown to promote clearance of hyperpigmented GBS from the female genital tract [37]. Further, previous studies have shown increased GBS vaginal persistence in B-cell-deficient mice or those lacking neonatal Fc-receptor, indicating that mucosal B cells and humoral immunity are important for GBS vaginal clearance [38]. It is possible that altered B cell numbers may contribute to the greater susceptibility of the Vɣ4/6^-/-^ mice, as it has been previously shown that they have reduced numbers of peripheral B cells [39]. Our study did not assess B cell or antibody responses to GBS over a time-course of colonization, but this would be of interest in the future, particularly in the context of recurrent colonization. While our study is the first to our knowledge to profile innate and T cell responses over a comprehensive time course of GBS vaginal colonization, other previous studies have also investigated immune responses to GBS in the FGT. For example, in a murine model of chronic GBS FGT colonization, lymphocyte and PMN cell infiltration were observed at the vaginal mucosal surface [17]. Interestingly, while cytokine responses were observed in vaginal tissues, this study found that changes in neutrophil and macrophage numbers were not observed within the submucosa by flow cytometry and GBS was not readily cleared from the FGT at these timepoints. Our laboratory has previously identified neutrophils in FGT tissue and has detected myeloperoxidase (MPO, used as an indicator of neutrophil recruitment) during colonization by hyperpigmented GBS [16]. In the present study, we also observed neutrophils in FGT tissues during GBS colonization by flow cytometry; however, we only observed slight, non-significant GBS-dependent increases in neutrophil numbers in the tissues at some early timepoints. Interestingly, induction of cytokine responses in the absence of robust immune infiltrate was also observed in murine model of GBS urinary tract infection, indicating GBS may broadly avoid immune infiltration during mucosal colonization/infection [40]. It is possible that while we did not observe differences in immune cell numbers, there may be functional or phenotypic immune changes during GBS colonization that we did not capture with our flow cytometric panels. Indeed, depletion experiments in this study indicated that FGT neutrophils may be important for controlling GBS colonization and uterine ascension. This was similarly reported in a recent study in pregnant mice, which observed increased GBS burden upon neutrophil depletion in the vagina and placenta (although not the cervix and uterus) [41].

Our results suggest that a steady immune state may facilitate persistent GBS colonization in the cervicovaginal tract over time. In a longitudinal study tracking women’s colonization status, 60% of the GBS-colonized cohort were persistently colonized throughout their pregnancies, often with the same isolate of GBS, with some considered “chronically” colonized since they tested GBS-positive at each sampling over the 21 month study period [42]. Further, in a study assessing GBS colonization pre- and post-pregnancy (∼8-10 weeks apart), almost 60% of women were colonized at both timepoints, and in 82% of these women the sequence type/serotype of the isolate did not change [43]. Further, several studies have observed that more than one third of women had recurrent GBS colonization in a subsequent pregnancy [44]. One interpretation of these data is that mounting of a successful, clearing anti-GBS immune response may be inconsistent with the ability of GBS to colonize long-term and recurrently. Collectively, these data are consistent with GBS being a persistent, albeit sometimes intermittent, commensal within the cervicovaginal tract and our results may help to explain the long-term and recurrent nature of GBS colonization in women. In contrast, during pregnancy, immune infiltration is characteristic of chorioamnionitis [45], and studies have modeled this inflammation and infiltration within gestational tissues during GBS infection of pregnant mice that mimics human disease [18, 46, 47]. While GBS has been shown to induce neutrophil and placental macrophage extracellular traps (NETs, METs) *ex vivo* [46, 48] and results in exacerbated inflammation in gestational membranes, increasing reports suggest that GBS may also thwart host defense, including subversion of killing by NETS [18, 49–51]. A recent study also showed that maternal immunity may be dampened during gestational diabetes, thereby promoting GBS perinatal infection [41]. It is possible that GBS may elicit increased immune infiltration at the maternofetal interface compared to the cervicovaginal tract and that alterations in hormones during pregnancy and/or in maternal metabolism in diabetes may further change FGT immune responses.

IL-17 responses constitute host defense against extracellular bacteria. Briefly, IL-23 production by epithelium or innate immune cells following pathogen detection stimulates mucosal T cells to produce IL-17. This induces epithelial upregulation of antimicrobial peptides as well as chemokines, which recruits neutrophils to the site of infection [52]. Previous work from our laboratory found that IL-17 levels are increased in vaginal tissue of GBS-colonized mice compared to PBS-colonized controls and that IL-17^+^ populations were associated with mice that had cleared GBS colonization [23]. IL-17 is produced by a variety of cell types including ɑβ T cells (Th17) [53], ɣδ T cells [54], CD8 T cells [55], mucosal associated invariant T (MAIT) cells [56], invariant Natural Killer T (iNKT) cells [57–59], as well as some innate populations, including ILC3s [60], alveolar macrophages [61] and neutrophils [62]. ɣδ T cells expressing the Vɣ6/Vδ1+invariant TCR [37] represent one of the first T cell types to colonize mucosal sites early in development [36] and are primary early producers of IL-17 in many disease and infection settings, characterized by their quick induction by IL-23 (independent of TCR stimulation) and importance in controlling infection [33, 54]. Similar to previous studies [63, 64], we show here that naïve FGT ɣδ T cells from CD-1 outbred mice are capable of producing IL-17 following stimulation. While we did not observe an expansion of total γδ T-cells in the FGT upon GBS colonization, our primary T cell panel did not include examination of those expressing Vɣ6. Therefore, we were unable to determine whether numbers of Vɣ6 ɣδ T cells specifically changed during GBS vaginal colonization. Despite this, IL-17 producing ɣδ T cells appear to be important for controlling GBS colonization as evidenced by increased GBS burdens in FGT tissues of Vɣ4/6^-/-^ mice compared to C57BL/6J WT mice. Future work will assess the contribution of IL-17 from other sources, such as Th17, MAIT, and iNKT cells and its impact on GBS vaginal colonization. These additional innate T cell populations may be of particular relevance as GBS encodes riboflavin biosynthesis machinery and therefore likely produces the 5-OP-RU ligand recognized by MAIT cells [65] and as iNKT cells have been previously shown to recognize group B streptococcal diacylglycerol-containing glycolipids [66] and to mediate immune responses upon immunization with a synthetic non-toxic derivative of the GBS toxin [67].

Altogether, this work represents a step towards understanding the complex interplay between GBS and the host immune systems during vaginal colonization and persistence in the FGT. GBS employs many immune evasive techniques, nicely reviewed elsewhere [68, 69], including the expression of the the β-hemolysin/cytolysin, a hemolytic pigment that promotes killing of a variety of immune cells [70, 71, 49, 72]. Recently, our laboratory has also identified a type VII secretion system in GBS that promotes vaginal persistence [22], and our current work is investigating a role for the T7SS effectors in modulating immune responses to GBS in the FGT. Understanding the host response to GBS during FGT colonization may facilitate new methods to limit GBS persistence in this niche as well as vertical transmission during pregnancy.

## Statements

### Acknowledgement (optional)

We acknowledge Drs. Harsha Krovi, and Henri Jupille, in technical development and consulting on these protocols and flow cytometric panels. We would also like to thank Jeremy Rahkola for his assistance with optimization of the flow cytometric panel and analysis and Arianne Crossen for technical assistance.

### Statement of Ethics

Animal experiments were performed using accepted veterinary standards as approved by the Institutional Animal Care and Use Committee at University of Colorado-Anschutz protocol #00316. The University of Colorado-Anschutz is AAALAC accredited, and their facilities meet and adhere to the standards in the “Guide for the Care and Use of Laboratory Animals”.

### Conflict of Interest Statement

The authors have no conflicts of interest to declare.

### Funding Sources

Funding for this work was provided by NIH/NIAID T32 AI007405 and NIH/NIAID F32 AI143203 to BLS, NIH/NIAID F32 AI186285 to SMM, and NIH/NIAID R01 AI153332 and NIH/NINDS R01 NS116716 to KSD.

### Author Contributions

Brady Spencer: Conceptualization, Investigation, Formal Analysis, Methodology, Visualization, Writing— Original draft and Review/Editing. Dustin Nguyen: Investigation, Formal Analysis, Methodology, Visualization, Writing—Original draft and Review/Editing. Stephanie Marroquin: Investigation, Review/Editing. Laurent Gapin: Resources, Methodology, Writing—Review/Editing. Rebecca O’Brien: Resources, Methodology, Writing—Review/Editing. Kelly Doran: Conceptualization, Funding Acquisition, Supervision, Writing— Review/Editing.

### Data Availability Statement

The data that support the findings of this study are available from the corresponding author upon reasonable request.

## Notes

### Competing Interest Statement

The authors have declared no competing interest.

