## Supplemental Material for "Characterization of the Cellular Immune Response to Group B Streptococcal Vaginal Colonization"

**Running title:** Cellular FGT immune responses to GBS

| Innate Panel | | | | | |
|---|---|---|---|---|---|
| **Marker** | **Fluorophore** | **Clone** | **Staining Concentration (μg/mL)** | **Company** | **Catalog #** |
| Viability Dye | eF506 | -- | [1:800 dilution] | Invitrogen | 65-0866-18 |
| CD45 | BUV395 | 30-F11 | 0.50 | BD Biosciences | 564279 |
| Ly6G | APC | 1A8 | 0.50 | Invitrogen | 17-9668-82 |
| CD11b | FITC | M1/70 | 2.50 | Invitrogen | 11-0112-85 |
| CD11c | PE-Cy7 | N418 | 1.00 | Biolegend | 117317 |
| CD64 | BV650 | X54-5/7.1 | 4.00 | BD Biosciences | 740622 |
| F4/80 | BV785 | BM8 | 1.00 | Biolegend | 123141 |
| Ly6C | BV421 | AL-21 | 0.50 | BD Biosciences | 562727 |
| I-A/I-E | BUV737 | M5/114 15.2 | 0.10 | BD Biosciences | 748845 |
| CD24 | PerCP-Cy5.5 | M1/69 | 0.20 | Biolegend | 101824 |
| CD49b | PE | DX5 | 4.00 | Biolegend | 108908 |
| RORγt | PE-CF594 | Q31-378 | 0.50 | BD Biosciences | 562684 |

| T Cell Panel | | | | | |
|---|---|---|---|---|---|
| **Marker** | **Fluorophore** | **Clone** | **Staining Concentration (μg/mL)** | **Company** | **Catalog #** |
| Viability Dye | eF506 | -- | [1:800 dilution] | Invitrogen | 65-0866-18 |
| CD45 | BUV395 | 30-F11 | 0.50 | BD Biosciences | 564279 |
| I-A/I-E | PE | M5/114 15.2 | 0.10 | Biolegend | 107608 |
| TCRγδ | BV650 | GL3 | 0.50 | BD Bioscience | 563993 |
| TCRβ | BV421 | H57-597 | 0.50 | BD Biosciences | 562839 |
| CD8 | BUV737 | 53-6.7 | 0.50 | BD Biosciences | 612759 |
| CD4 | FITC | H129.19 | 1.25 | Biolegend | 130308 |
| CD69 | BV785 | H1.2F3 | 1.00 | Biolegend | 104543 |
| CD44 | PerCP-Cy5.5 | IM7 | 1.00 | Biolegend | 103032 |
| Foxp3 | APC | FJK-16S | 2.00 | Invitrogen | 17-5773-82 |
| CD122 | PE-Cy7 | TM-β1 | 2.00 | Biolegend | 123216 |
| RORγt | PE-CF594 | Q31-378 | 0.50 | BD Biosciences | 562684 |

**Supplemental Table 1. Innate and T cell flow cytometric panels used in the present study.**

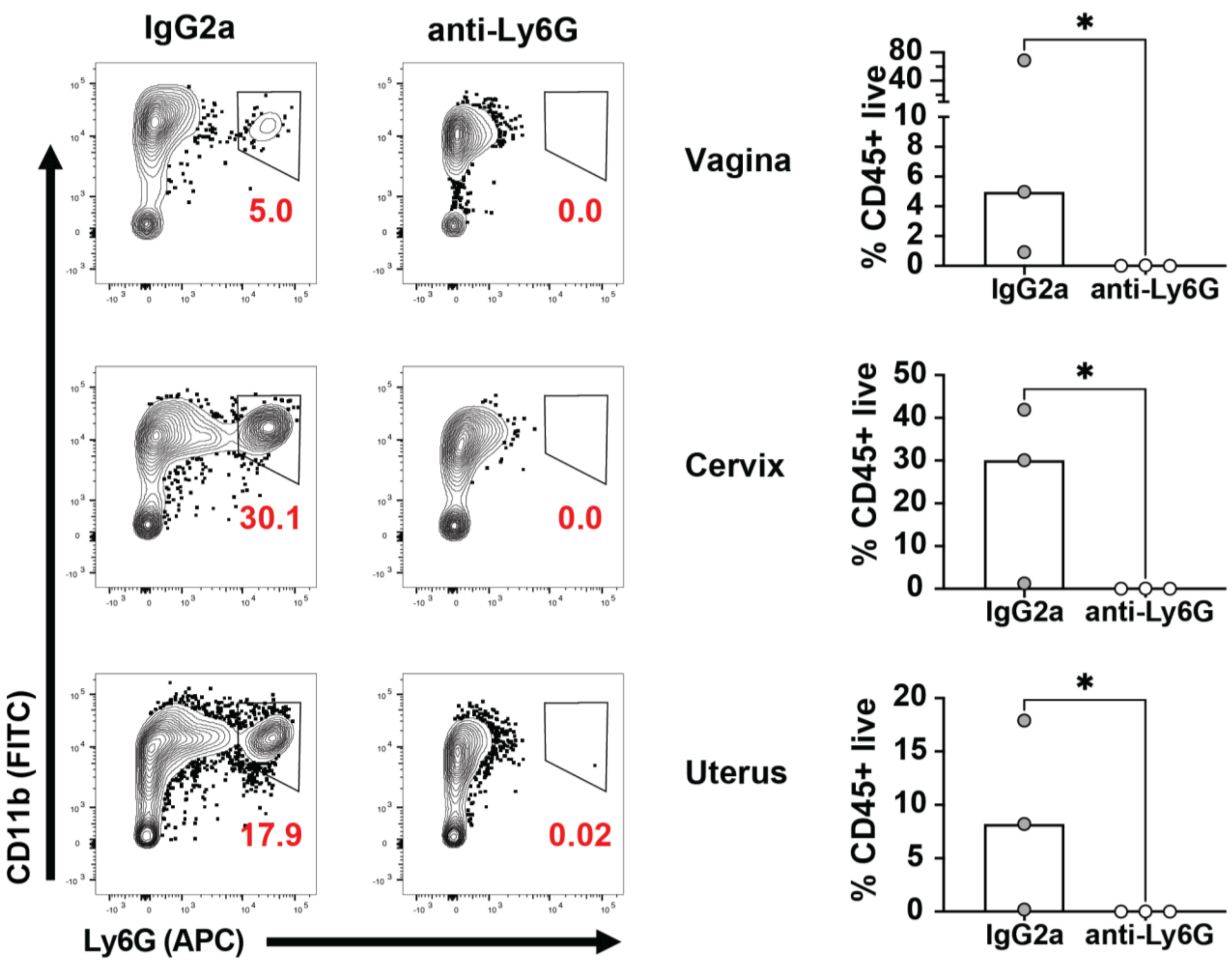


**SFig. 1. Validation of neutrophil depletion in FGT tissues and impact on COH1 vaginal colonization.** Cd11b+ Ly6G+ double positive neutrophils were evaluated in the vagina, cervix, and uterus of mice receiving IgG2a isotype control or anti-Ly6G depleting antibody as displayed in Fig 1a. Flow cytometry confirmed a lack of neutrophils in anti-Ly6G treated animals at day 4 post-PBS inoculation. Red text indicates neutrophils percentages (of live $CD45^{+}$ immune cells). Neutrophil percentages of n = 3 mice/group are shown on the right for each tissue. Bars indicate the median and statistics represent a one-tailed Mann Whitney U test.

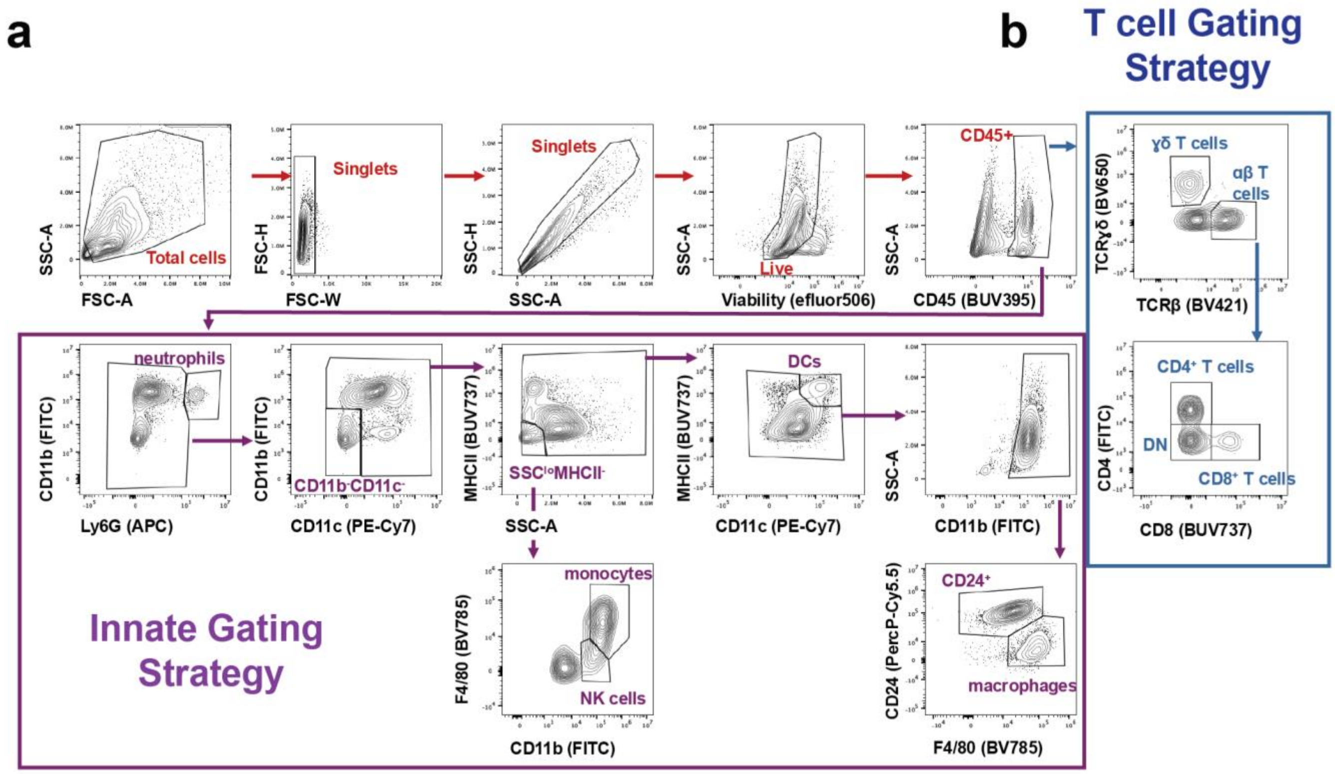


**SFig 2. Gating strategy of innate and T cell populations in the murine FGT.** Contour plots of a murine uterine single cell suspension stained with an innate antibody panel or a T cell antibody panel. Gates identifying cell populations of interest are denoted in red, blue or purple text. **a)** The innate panel identifies neutrophil, $Cd11b^-$ $CD11c^-$, $SSc^{lo}$/$MHCII^-$, dendritic cell, macrophage, and $CD24^+$ populations. **b)** The T cell panel identifies ɣδ T cells as well as αβ T cells, including $CD4^+$ and $CD8^+$ and double negative αβ T cell populations. The gating strategy shown is representative of 3 independent flow time-course experiments and of vaginal, cervical, and uterine tissue.

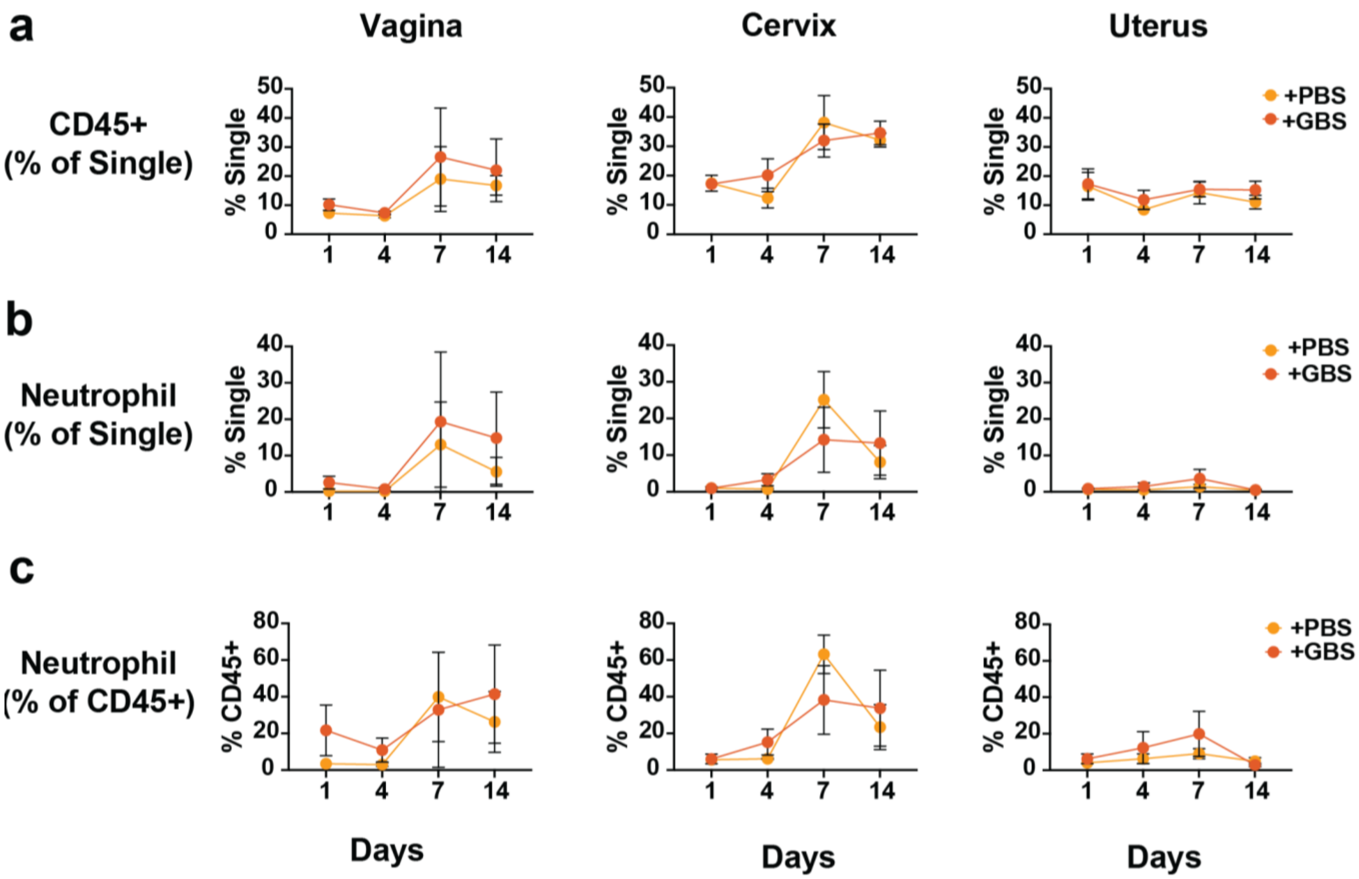


**SFig 3. Total immune cells and neutrophils are not significantly altered over a time-course of COH1 vaginal colonization.** Line graphs depict **a**) CD45$^+$ immune cells as a proportion of single cells and **b**) neutrophils (as a proportion of single cells) or **c**) as a proportion of CD45$^+$ immune cells over a time-course of GBS colonization at days 1, 4, 7, and 14. Yellow lines indicate PBS mock-colonized mice and orange lines indicate GBS-colonized mice. Each data point represents the mean and error bars indicate standard deviation. The data represent a total of n = 3 per timepoint. Statistics represent two-way ANOVA with Šídák's multiple comparisons test.

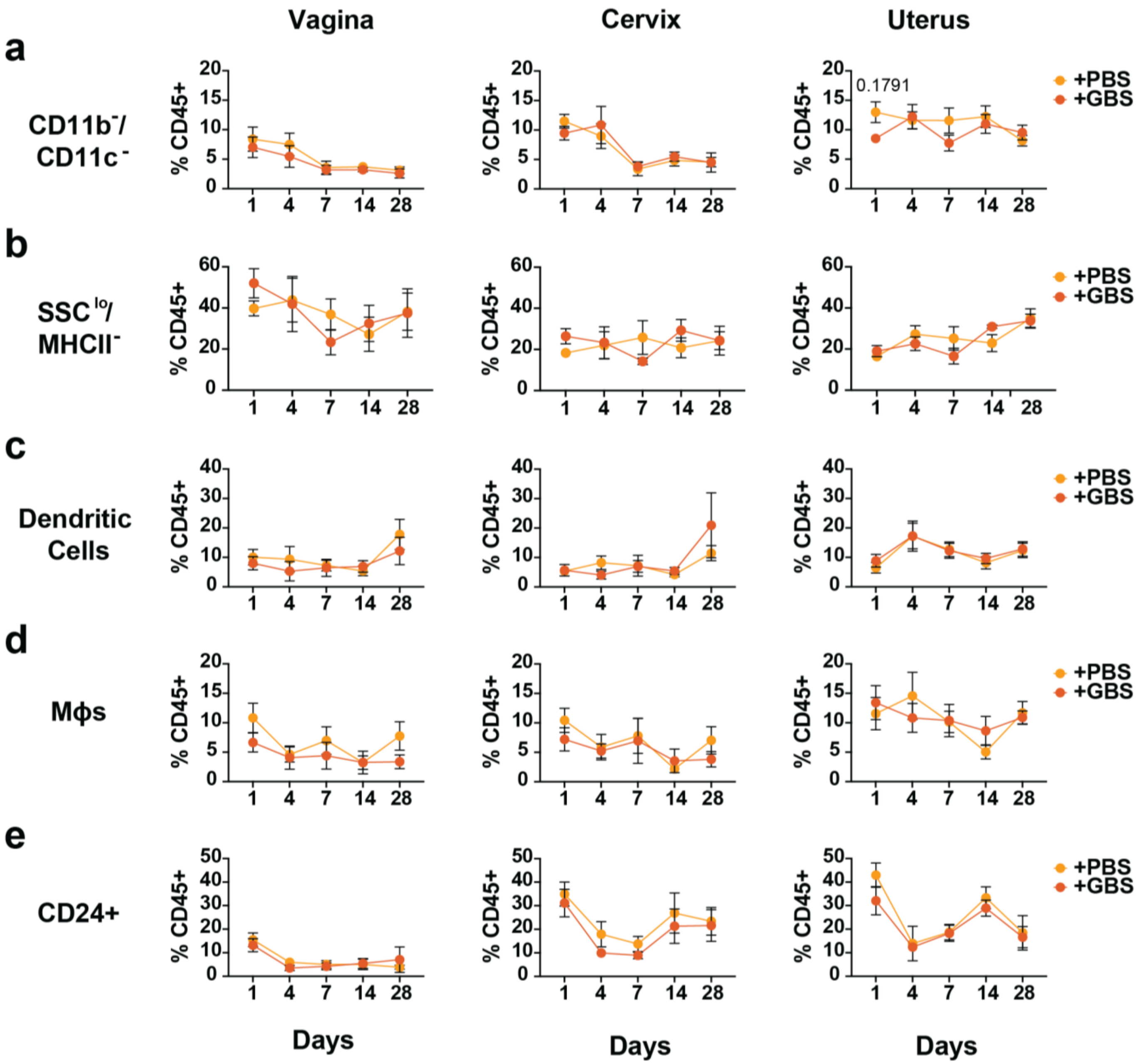


**SFig 4. Other innate immune populations are not significantly altered over a time-course of CJB111 vaginal colonization.** Line graphs depict **a**) Cd11b$^-$CD11c$^-$, **b**) SSc$^{lo}$/MHCII$^-$, **c**) dendritic cell, **d**) macrophage, and **e**) CD24$^+$ populations as a proportion of CD45$^+$ immune cells over a time-course of CJB111 colonization at days 1, 4, 7, 14, and 28. Yellow lines indicate PBS mock-colonized mice and orange lines indicate CJB111-colonized mice. Each data point represents the mean and error bars indicate standard deviation. The data represent a total of n = 6-8 per timepoint. Statistics represent two-way ANOVA with Šídák's multiple comparisons test.

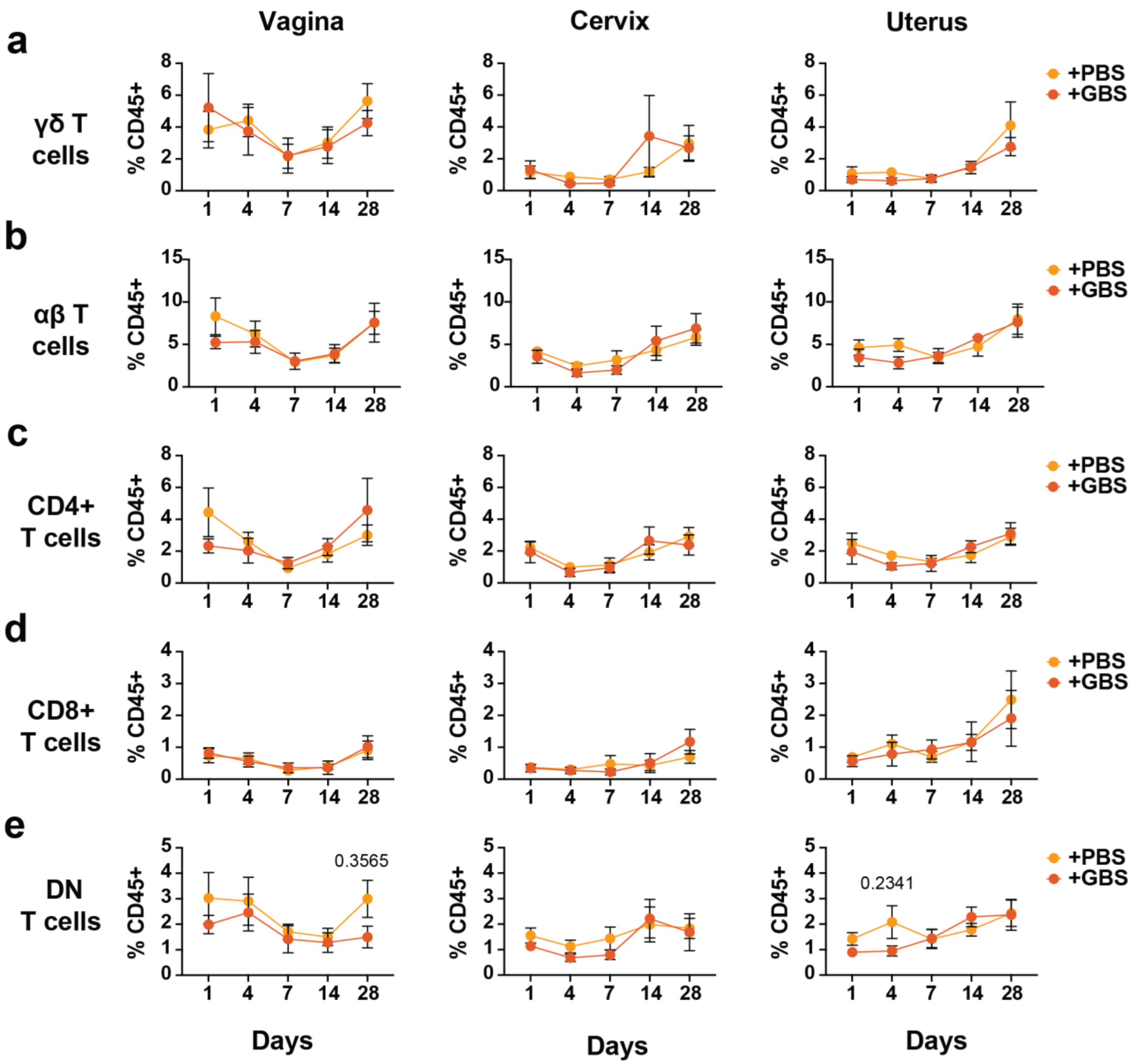


**SFig 5. T cell populations are not significantly altered over a time-course of CJB111 vaginal colonization.** Line graphs depict **a**) γδ T cells, **b**) αβ T cells, including **c**) $CD4^+$, **d**) $CD8^+$, and **e**) double negative αβ T cell populations as a proportion of $CD45^+$ immune cells over a time-course of CJB111 colonization at days 1, 4, 7, 14, and 28. Yellow lines indicate PBS mock-colonized mice and orange lines indicate CJB111-colonized mice. Each data point represents the mean and error bars indicate standard deviation. The data represent a total of n = 6-8 per timepoint. Statistics represent two-way ANOVA with Šídák's multiple comparisons test.
